# An open-access EEG dataset for speech decoding: Exploring the role of articulation and coarticulation

**DOI:** 10.1101/2022.11.15.516461

**Authors:** João Pedro Carvalho Moreira, Vinícius Rezende Carvalho, Eduardo Mazoni Andrade Marçal Mendes, Ariah Fallah, Terrence J. Sejnowski, Claudia Lainscsek, Lindy Comstock

## Abstract

Electroencephalography (EEG) holds promise for brain-computer interface (BCI) devices as a non-invasive measure of neural activity. With increased attention to EEG-based BCI systems, publicly available datasets that can represent the complex tasks required for naturalistic speech decoding are necessary to establish a common standard of performance within the BCI community. Effective solutions must overcome various kinds of noise in the EEG signal and remain reliable across sessions and subjects without overfitting to a specific dataset or task. We present two validated datasets (N=8 and N=16) for classification at the phoneme and word level and by the articulatory properties of phonemes. EEG signals were recorded from 64 channels while subjects listened to and repeated six consonants and five vowels. Individual phonemes were combined in different phonetic environments to produce coarticulated variation in forty consonant-vowel pairs, twenty real words, and twenty pseudowords. Phoneme pairs and words were presented during a control condition and during transcranial magnetic stimulation targeted to inhibit or augment the EEG signal associated with specific articulatory processes.

## Background & Summary

Brain-computer interface (BCI) devices aim to restore communicative abilities to individuals who have lost motor function. BCI speech decoding devices must balance the need for a user-friendly interface with task performance^1^. Neural signals from electrocorticography (ECoG) can provide a measure of task-relevant brain activity with a minimal processing latency and a superior signal-to-noise ratio^2,3^. However, ECoG studies must be conducted on surgery patients who differ in many respects from the end user of BCI devices. Alternatively, electroencephalography (EEG) allows for BCI devices to be tested directly among the target population. But because neural signals become distorted as they pass through the skull and scalp to reach EEG sensors^4^, BCI applications are often limited to the detection of stimulus-evoked potentials^5–8^, resulting in non-naturalistic paradigms that remain comparatively slow and inflexible^9,10^. A practical solution to speech decoding will require the quick and accurate classification of a large inventory of context-dependent speech sounds in rapid succession^11^. To utilize EEG effectively for these purposes, researchers will benefit from datasets that can train models to systematically overcome the difficulties presented by the nature of the EEG signal and the complexity of the task.

Methodological paradigms for speech decoding include covert speech and auditory comprehension tasks^12–24^. The resulting analyses are heterogeneous in nature and few authors make their data publicly available, complicating efforts to reproduce and compare different decoding methods. The publication of EEG datasets allows researchers to benchmark model performance^25,26^. Currently available datasets typically focus on a task of high difficulty (e.g., imagined or inner speech) yet remain minimal in scope, with a small number of trials, stimulus types, and subjects. Moreover, there is little coherence between different stimulus types that would allow researchers to test their models against progressively more naturalistic data. BCI devices rely upon the neural signals associated with muscle movements^27^, and although the utility of classification schemes based on speech articulation has been noted by numerous researchers^28–32^, this factor has yet to be included as an organizing principle in published datasets. Finally, the overfitting of highly complex machine learning models is frequently discussed in the literature^33^ but rarely tested overtly. Independently collected datasets of the same data types are needed to ensure that models can generalize across subjects and sessions. We present two datasets for EEG speech decoding that address these limitations:

- Naturalistic speech is not comprised of isolated speech sounds^34^. The phonetic environment surrounding phonemes affects their quality^35,36^, complicating accurate category designation^37,38^. Existing datasets lack a wide range of co-articulated phonemes. We provide single, double, and phoneme triplets for six consonants (/b/, /p/, /d/, /t/, /s/, /z/) and five vowels (/i/, /E/, /A/, /u/, /oU/) to assess classification accuracy in progressively more complex phonetic environments.
- The decoding literature devotes considerable attention to the articulatory properties of phonemes^28–32^. The most successful decoding models integrate this knowledge^19,39,40^. Available datasets do not select phonemes that fall within overlapping articulatory categories (place, manner, voicing). We provide consonants that represent unique combinations of features (bilabial/alveolar; stop/fricative; voiced/unvoiced) to assess articulatory features as a classification parameter.
- The integration of probabilistic language models into BCI devices has predisposed researchers to anticipate better results when training on real words^41–44^. Published EEG datasets comprise real-word stimuli. Yet functional magnetic resonance imaging (fMRI) studies reveal a greater hemodynamic response to pseudowords^45,46^. Effortful processing may strengthen neural signals and facilitate decoding^47^. We provide twenty real and twenty pseudowords to test this hypothesis.
- Robust speech decoding models must compensate for a high degree of noise from various sources. In addition to the limitations of EEG data (noise from within the signal), sounds are often heard inaccurately in the presence of environmental noise or when the participant is inattentive to stimuli^48–50^. We provide data from a control condition and two TMS conditions that facilitate or impede task performance, depending on the phoneme type and stimulation target^51^.
- Speech decoding papers typically publish one or more analyses conducted on a single dataset. This raises concerns about overfitting and how well the model might perform on additional data without significant modification^52^. We provide two datasets (N=8 and N=16) collected at different time points for the same stimulus types and/or participants. The second serves as an external validation set to allow researchers to assess model performance on independently derived data^53^.

## Methods

Our aim was to create a systematically structured, multifaceted dataset in two parts that would represent a novel contribution to the publicly available data for EEG speech decoding. This involved a careful selection of stimulus types to ensure a scaffolded progression in the kind of classification analyses that could be conducted in terms of the choice of parameter and the complexity of the task (single phonemes, phoneme pairs, phoneme triplets). Some trials (single phonemes) represent neural data during comprehension and production. Other trials (phoneme pairs and triplets) take advantage of an experimental paradigm designed to either facilitate speech decoding or introduce neural noise. The outcome depends on whether the stimulation site is matched to the phoneme type. This paradigm improved subject performance in phoneme discrimination (in matched trials) by administering two closely spaced TMS pulses prior to phoneme presentation^54^. There is evidence that TMS may facilitate speech decoding when stimulation to the motor cortex target regions that control the muscles involved in the production of a specific phoneme category, and errors in decoding may be induced when stimulation occurs in an unrelated region of the motor cortex, prompting the perception of another phoneme category^51,54^. A mismatch between the phoneme type and stimulation site will introduce uncertainty into the decoding task, similar to when a phoneme is misheard in everyday speech. By collecting the same data type (CV phoneme pairs) at two separate time points, we invite researchers to develop their model on the larger dataset and then to assess the robustness of the model in a second, smaller and noisier validation dataset.

### Participants

Participants aged between 20 and 40 were recruited from the UCLA campus by means of flyers. Ten participants (6 female) were recruited for the first dataset in 2019 and twenty participants (10 female) were recruited for the second dataset in 2021. Inclusion criteria were defined as no diagnosis of any neurological, psychiatric, or developmental disorders, self-reported normal hearing, and no contraindications for TMS or MRI procedures (implanted medical devices, implanted metal, pregnancy, personal or family history of seizures, and exclusionary medications). Screening included the completion of an abbreviated version of the experimental task to ensure that participants understood the task directions and could perform the phoneme discrimination task. Left-hemisphere lateralization of the language processing regions in all participants was established during an fMRI scan in which participants performed the task. Two individuals (1 female) were excluded from the 2019 dataset due to necessary modifications made to the stimulus audio files after their participation. In the 2021 dataset, one participant (male) was excluded due to poor task performance and three participants (male) were excluded due to complications with the TMS equipment that may have led to imprecise targeting. Participants provided informed consent and were paid for two sessions. The experimental protocol was approved by the UCLA Institutional Review Board (IRB#21-000333).

### Experimental design

The study was conducted in three sessions (Fig. 1). In the first session, participants underwent a directed interview to ensure that they met the study inclusion criteria and possessed no contraindications. Participants who were able to perform an abbreviated phoneme perception task with at least 75% accuracy were enrolled. In the second session, participants underwent an MRI scan to aid in neuronavigation for the TMS procedure and performed the phoneme discrimination task in the MRI scanner to lateralize their language processing areas. The final session involved the recording of EEG signals while participants performed the phoneme perception task. TMS was targeted to areas of the motor cortex associated with the production of specific phonemes. In the second round of data collection that was conducted in 2021, participants also listened to and repeated single phonemes and performed the perception task for phoneme triplets.

**Figure 1.**
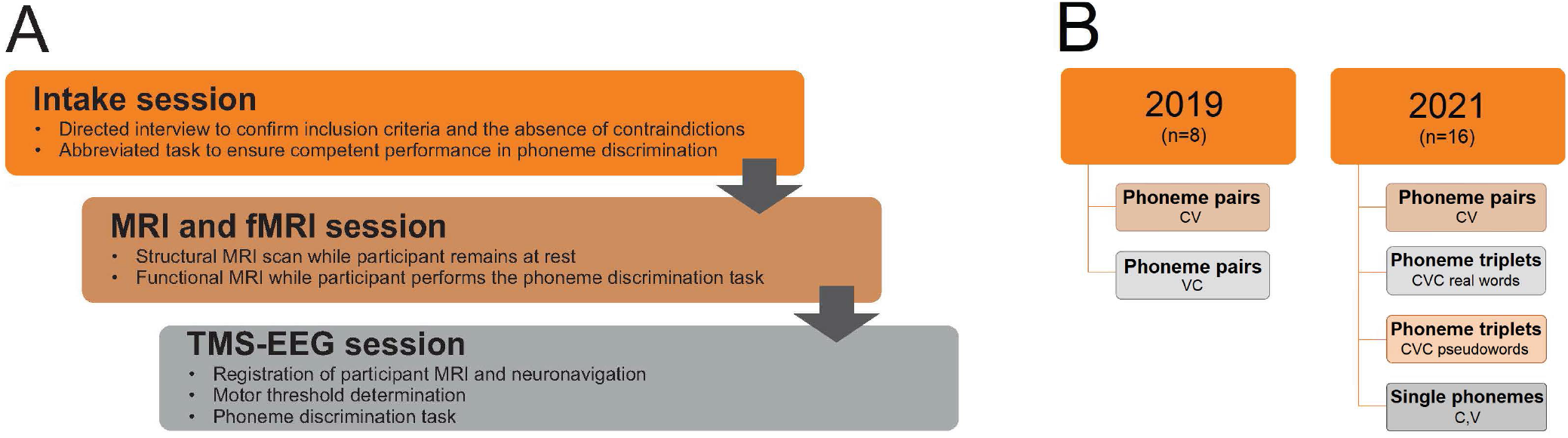
Session organization. (A) The experiment was conducted in three sessions that were held on separate days. Participant eligibility was confirmed in the first session. MRI scanning and TMS-EEG data collection were conducted independently. (B) Additional participants and trial types were included in 2021.

### Data collection

#### MRI scanning

Scanning was conducted in the UCLA Center for Cognitive Neuroscience with a Siemens Prisma-FIT 3T Scanner. Participants were provided with ear protectors and headphones for a 45 to 60 dB reduction of the noise associated with scanning, thus ensuring that participants could hear the stimuli clearly and that the noise level was not uncomfortably loud. Participants were asked to lie with their head motionless during all scanning procedures. High-resolution anatomical images were acquired, followed by a functional scan in which participants were directed to either relax passively while looking at a fixation cross or to perform the button-press phoneme discrimination task. Functional data was acquired in a block design with a BOLD-weighted echoplanar imaging sequence aligned in parallel to the bicommissural plane, yielding 36 slices covering the whole brain, each 3 mm thick with a 1 mm gap between slices. Each slice was acquired as a 64 × 64 matrix yielding an in-plane resolution of 1.5 × 1.5 mm. The total duration of the scanning session was 40 minutes.

#### TMS-EEG

The TMS-EEG procedure was conducted in the Neuromodulation Division of the Semel Institute for Neuroscience and Human Behavior at UCLA. The TMS equipment included a Magstim Super Rapid Plus1 stimulator and a figure-of-eight 40 mm coil. The EEG system included an eego™ sports WaveGuard 64-channel EEG cap and eego mylab system compatible with electromagnetic stimulation. Targeting was completed using the Visor 2 neuronavigation system. The electrode positions were digitized and registered to individual participant MRIs using the ANT Neuro Xensor. EEG signals were bandpass-filtered 0.1-350 Hz, sampled at 2000 Hz, and referenced to the CPz electrode. All electrode impedances were kept < 5 kΩ. Stimulus presentation and recording of reaction time data were collected using PsychoPy^55^.

The appropriate stimulation intensity for TMS studies is determined on an individual basis^56^. Prior to the experimental session, the motor threshold (rMT) of each participant was determined by eliciting motor-evoked potentials (MEPs) in the first dorsal interosseus (FDI) muscle of the dominant hand at the minimum amount of stimulation needed to evoke an MEP in a hand muscle after a single pulse over M1. Single TMS pulses were delivered to locations in the motor cortex contralateral to the dominant hand. The intensity of the stimulation was gradually lowered until reaching a level of stimulator output at which 5 out of 10 MEPs in the hand muscle had an amplitude of at least 50 microvolts. Potentials evoked during TMS represent the net sum of excitatory and inhibitory stimulation effects^57–59^. The literature has found that excitation increases at intensities of 110-120% rMT. In accordance with our reference study, stimulation was administered at 110% of the FDR rMT^54^. A physician observed the motor thresholding procedure to ensure that no negative effects were incurred by participants.

TMS targeted areas of the motor cortex involved in (i) lip and tongue movements (which produce bilabial and alveolar consonants, respectively) or (ii) processing of real and pseudowords. Stimulation targets were defined as the MNI coordinates of peak motor cortex activation in LipM1 and TongueM1 during lip and tongue articulatory movements (lips: −56, −8, 46; tongue: −60, −10, 25), taken from the literature^60^ and the reference study^54^. However, cortical functional localization shows considerable individual variation^61^. Therefore, the coordinates were assessed relative to the fMRI task results of each participant to ensure overlap between the targets and individual task localization. The second set of targets was defined after observing common regions of fMRI activation across participants during the relevant blocks of the fMRI task. Broca’s area (BA 44: −51, 7, 23) was the target for real words and a region implicated in verbal memory (BA 6: −46, 1, 41) was the target for pseudowords^62^.

#### Behavioral task

The phoneme discrimination task consisted of listening to speech sounds and identifying stimuli with a button-press response. Auditory stimuli (Fig. 2) were presented via laptop speakers: (i) single phonemes, (ii) paired consonant-vowel phonemes (CV, VC), and (iii) real or pseudowords constructed of phoneme triplets (CVC). Consonant stimuli included four phonemes in the pair and triplet conditions (/b/, /p/, /d/, /t/), with the addition of two additional phonemes (/s/, /z/) in the single condition. Vowel stimuli included five phonemes in all conditions (/i/, /E/, /A/, /u/, /oU/). These sets yielded 11 individual phonemes (6 consonants and 5 vowels), 40 phoneme pairs (20 CV/20 VC), and 40 phoneme triplets (20 real/20 pseudowords). Participants were asked to listen to and repeat the phoneme in the single condition, to identify the consonant phoneme in paired conditions, and to identify phoneme triplets as real or pseudowords. Multiple classification analyses may be conducted on each stimulus type (Fig. 3).

**Figure 2.**
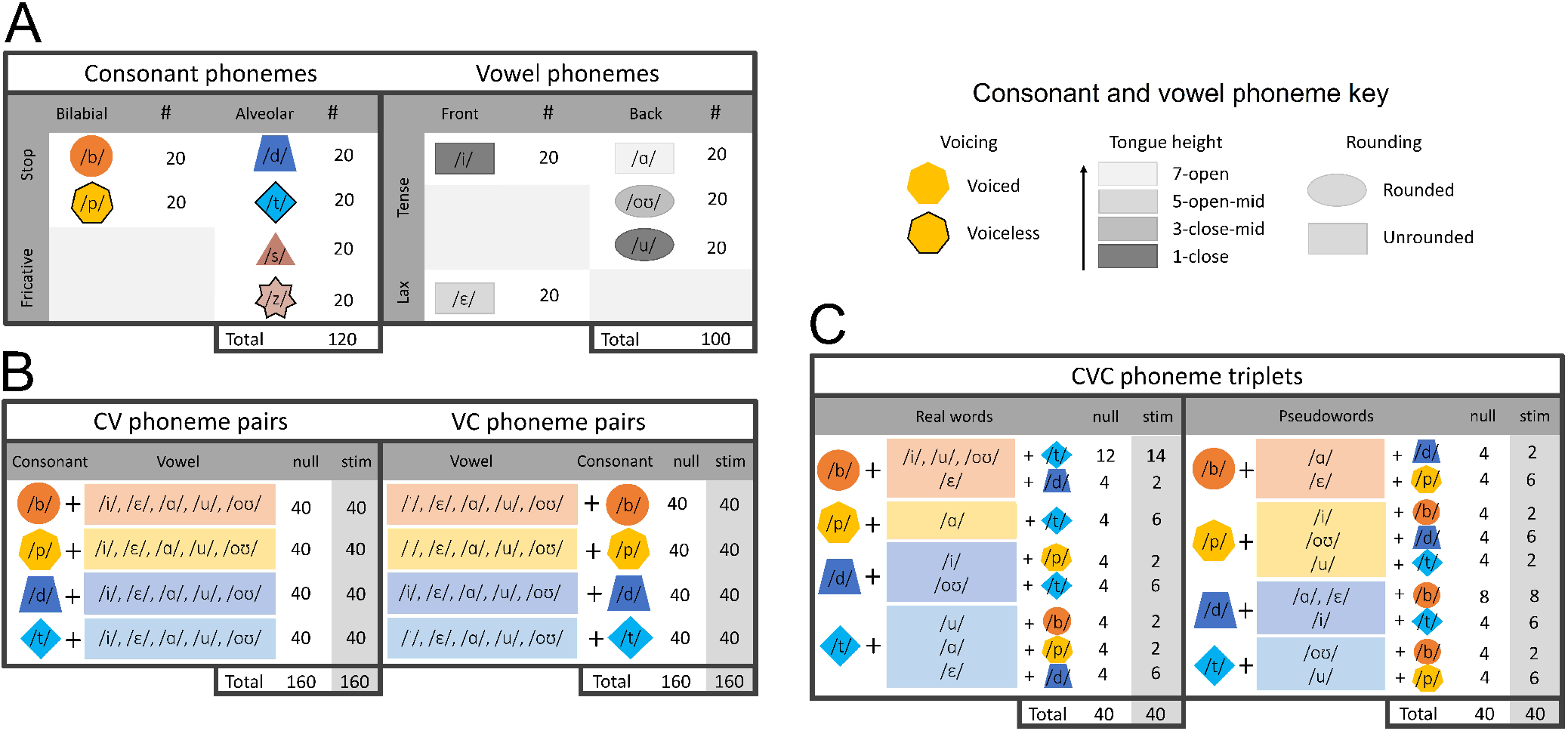
Stimulus types. (A) Consonant and vowel phonemes possess unique articulatory features. Consonants can be described by three parameters: place of articulation (bilabial, alveolar), manner of articulation (stop, fricative), and voicing (voiced, unvoiced). Vowels can be described by four parameters: tongue height (from close to open), tongue position (front, back), tongue tension (tense, lax), and lip position (rounded, unrounded). (B) Phoneme pairs included eight instances of each combination of stop consonants and vowels. (C) Phoneme triplets were limited by the stipulation to create real and pseudowords. Stimuli included eight instances of each vowel in the real and pseudoword conditions.

**Figure 3.**
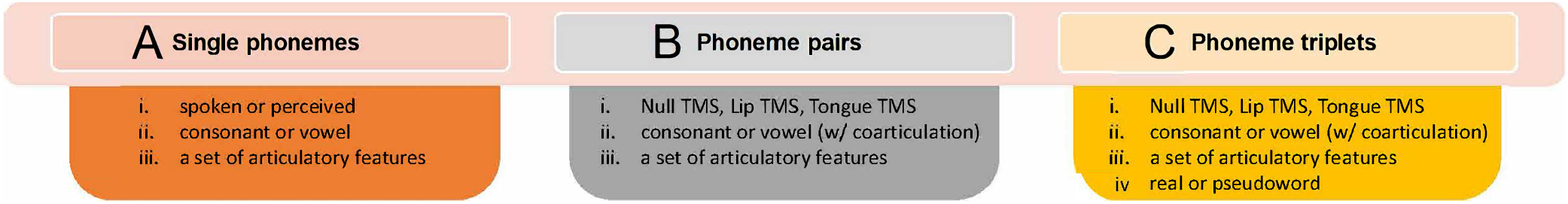
Analysis. (A) Single phonemes can be classified by modality, category, and articulation. (B) Phoneme pairs – by target, category, and articulation. (C) Phoneme triplets – by target, category, articulation, and word type.

TMS elicits a period of excitatory activation with an onset latency of 50-80 ms after stimulation^63^. We reproduced the design of our reference study^54^ to ensure an excitatory neural response that would translate into task facilitation. Each trial delivered paired TMS pulses at one of the stimulation targets, separated by a short interpulse interval (50 ms). Excitation of the cortical region not involved in stimulus production (i.e., TMS at LipM1 during alveolar phoneme presentation) results in neural noise that interferes with the perception task. The audio stimulus followed 50 ms after the second TMS pulse. One target was stimulated per run (counterbalanced across participants). Details of the experimental protocol are illustrated in Figure 4.

**Figure 4.**
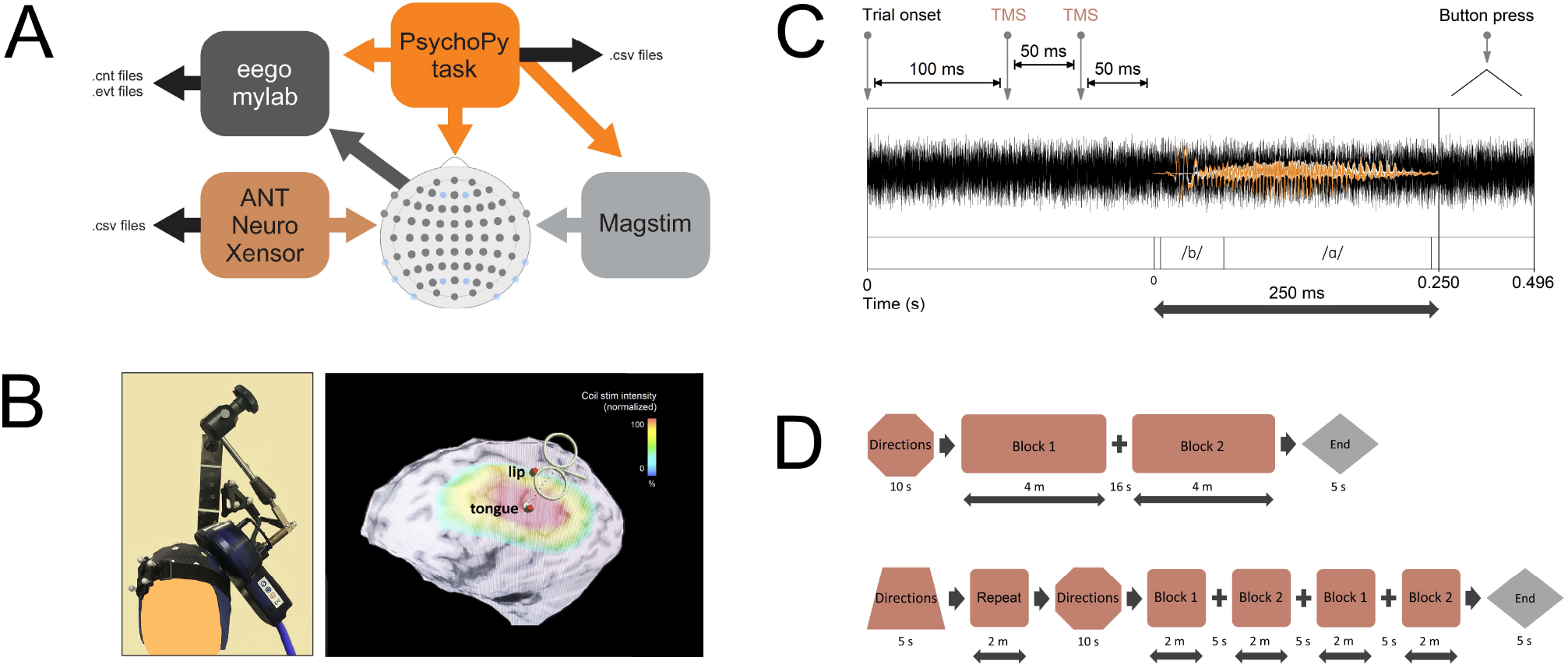
Experimental protocol. (A) Software controlled the experimental task and sent triggers to initiate TMS and to create timestamps for each pulse. (B) The TMS coil was positioned at the necessary stimulation site. Despite some overlap in the induced magnetic field, only the targeted region received maximum stimulation intensity. (C) The audio stimuli were immersed in white noise. Two TMS pulses were administered 50 ms prior to stimulus onset. (D) The run design in 2019 was reproduced in 2021 with the addition of more frequent breaks between blocks. A block of single phonemes was introduced in 2021.

Participants listened to audio clips immersed in 500 ms of white noise. The white noise created a mild background distraction for participants to ensure that they did not perform the phoneme discrimination task at ceiling. Participants were instructed to respond as fast as possible with a button press after they had identified the phoneme. In the case of multiple button presses, correct trials were determined from the initial button press. Participants who exhibited a non-random response strategy (i.e., failure to select from the full set of phonemes) were excluded. In 2021, participants were instructed to listen to single phonemes without TMS and to repeat the sound they heard immediately after stimulus presentation (300 ms from trial onset).

Two lists of stimulus items were used with one list assigned to each block. In 2019, the runs were split into two blocks. The first block presented CV pairs, followed by a block of VC pairs. In 2021, four blocks were administered per run. The first two blocks presented CV pairs, followed by two blocks of CVC stimuli (real and pseudowords). A five-minute break was provided between runs. Participants completed 120 trials in each run: 80 with TMS and 40 random catch trials. In 2021, each run of the task was preceded by the presentation of 220 trials of single phonemes (20 trials each). Stimuli in all conditions were presented in a pseudo-randomized order. The total run time of the experiment lasted 49 minutes in 2019 and 58 minutes in 2021.

Minimal modifications to this procedure were made during the intake and scanning sessions. For the initial assessment, half of the task was administered. During fMRI data collection, the full-length task was administered. However, stimuli were presented in blocks of the same type (bilabial, alveolar, real words, pseudowords) to aid in their cortical localization.

### Data characterization

#### Classification of acquired data

The procedure required sustained attention during a lengthy TMS procedure. The mean reaction time and standard deviation were calculated to confirm that participants were attentive to the task throughout the procedure. These metrics are documented in `.csv` files uploaded to the data repository. In the 2019 dataset, some variation in trial numbers is observed due to missed trials and rotation in the list of stimuli administered to each participant. No subjects performed less than 90% of the total list, with the exception of P04 in the VC condition with LipTMS. Here, excluded trials resulted from missed trials. In the 2021 dataset, all trials were uploaded irrespective of a button-press response. Two subjects performed an abbreviated list of phoneme triplets, and one also performed an abbreviated list of single phonemes. The number of tagged trials is shown in Table 1.

**Table 1.** Final number of trials presented by participant, target, and stimulus type. (A) 2019 dataset. (B) 2021 dataset.

| Task | Target | S01 | S02 | S03 | S04 | S05 | S06 | S07 | S08 | Total |
| --- | --- | --- | --- | --- | --- | --- | --- | --- | --- | --- |
| Lip TMS | CV | 79 | 81 | 80 | 76 | 77 | 80 | 81 | 77 | 631 |
|  | VC | 77 | 79 | 79 | 57 | 79 | 79 | 78 | 78 | 606 |
| Lip NULL | CV | 39 | 37 | 38 | 38 | 36 | 37 | 38 | 38 | 301 |
|  | VC | 41 | 43 | 42 | 33 | 41 | 41 | 42 | 42 | 325 |
| Tongue TMS | CV | 80 | 81 | 81 | 78 | 77 | 79 | 79 | 76 | 631 |
|  | VC | 80 | 78 | 78 | 78 | 77 | 79 | 78 | 76 | 624 |
| Tongue NULL | CV | 39 | 38 | 38 | 38 | 36 | 38 | 38 | 36 | 301 |
|  | VC | 41 | 42 | 41 | 41 | 38 | 41 | 40 | 38 | 322 |
| Total |  | 476 | 479 | 477 | 439 | 461 | 474 | 474 | 461 |  |

| Target | Task | S01 | S02 | S03 | S04 | S05 | S06 | S07 | S08 | S09 | S10 | S11 | S12 | S13 | S14 | S15 | S16 | Total |
| --- | --- | --- | --- | --- | --- | --- | --- | --- | --- | --- | --- | --- | --- | --- | --- | --- | --- | --- |
|  | Single spoken | 110 | 220 | 220 | 220 | 220 | 220 | 220 | 220 | 220 | 220 | 220 | 220 | 165 | 220 | 220 | 220 | 3355 |
|  | Single perceived | 110 | 220 | 220 | 220 | 220 | 220 | 220 | 220 | 220 | 220 | 220 | 220 | 165 | 220 | 165 | 165 | 3355 |
| Lip | CV pairs | 80 | 80 | 80 | 80 | 80 | 80 | 80 | 80 | 80 | 80 | 80 | 80 | 80 | 80 | 80 | 80 | 1280 |
| Lip NULL | CV pairs | 40 | 40 | 40 | 40 | 40 | 40 | 40 | 40 | 40 | 40 | 40 | 40 | 40 | 40 | 40 | 40 | 640 |
| Tongue | CV pairs | 80 | 80 | 80 | 80 | 80 | 80 | 80 | 80 | 80 | 80 | 80 | 80 | 80 | 80 | 80 | 80 | 1280 |
| Tongue NULL | CV pairs | 40 | 40 | 40 | 40 | 40 | 40 | 40 | 40 | 40 | 40 | 40 | 40 | 40 | 40 | 40 | 40 | 640 |
| BA 44 | Real words | 0 | 30 | 40 | 40 | 40 | 40 | 40 | 40 | 40 | 40 | 40 | 40 | 0 | 40 | 40 | 40 | 550 |
|  | Pseudowords | 0 | 31 | 40 | 40 | 40 | 40 | 40 | 40 | 40 | 40 | 40 | 40 | 0 | 40 | 40 | 40 | 551 |
| BA 44 NULL | Real words | 0 | 16 | 20 | 20 | 20 | 20 | 20 | 20 | 20 | 20 | 20 | 20 | 0 | 20 | 20 | 20 | 276 |
|  | Pseudowords | 0 | 16 | 20 | 20 | 20 | 20 | 20 | 20 | 20 | 20 | 20 | 20 | 0 | 20 | 20 | 20 | 276 |
| BA 6 | Real words | 40 | 40 | 40 | 40 | 40 | 40 | 40 | 40 | 40 | 40 | 40 | 40 | 80 | 40 | 40 | 40 | 680 |
|  | Pseudowords | 40 | 40 | 40 | 40 | 40 | 40 | 40 | 40 | 40 | 40 | 40 | 40 | 80 | 40 | 40 | 40 | 680 |
| BA 6 NULL | Real words | 20 | 20 | 20 | 20 | 20 | 20 | 20 | 20 | 20 | 20 | 20 | 20 | 40 | 20 | 20 | 20 | 340 |
|  | Pseudowords | 20 | 20 | 20 | 20 | 20 | 20 | 20 | 20 | 20 | 20 | 20 | 20 | 40 | 20 | 20 | 20 | 340 |
| Total |  | 690 | 893 | 920 | 920 | 920 | 920 | 920 | 920 | 920 | 920 | 920 | 920 | 810 | 920 | 920 | 920 |  |

### Data processing

The continuous raw data from the EEG recordings have been uploaded to the data repository so that researchers may apply their preferred pre-processing and processing pipeline, as necessary for alternative speech decoding models. However, in order to exemplify the principle of transparency, we have also made available three stages of data cleaning or signal processing that were utilized for our data validation section (i) data normalized within each data window to zero mean and unit variance (for DDA); (ii) data resampled at 256Hz, filtered (notch filter in the 59Hz to 61Hz bands and band-pass filter from 0.1Hz to 100Hz), and separated by trial (for ERPs); and (iii) data that underwent additional processing, containing two rounds of cleaning of unwanted ICA components in most cases (for ERPs).

A Matlab routine was designed based on the EEGLAB library^64^ for data analysis and pre-processing. The code (available in the Code Availability section) is separated into several sections, each responsible for a part of the data processing. This makes it possible to organize, streamline, and automate the analysis, in a process that eliminates the extensive use of EEGLAB interface by the user in cases where it is not strictly necessary. The routine is structured as shown in Figure 5 and consists of seven main sections that include the removal of unwanted channels, event setup based on the information tables, resampling and filtering, separation of trials based on events, visual inspection for cleaning up bad trials, ICA decomposition, and manual removal of unwanted ICA components. In addition, the code offers optional sections for interpolation of TMS signals, generation of signal state images, and other tools that provide information on changes made during processing.

**Figure 5.**
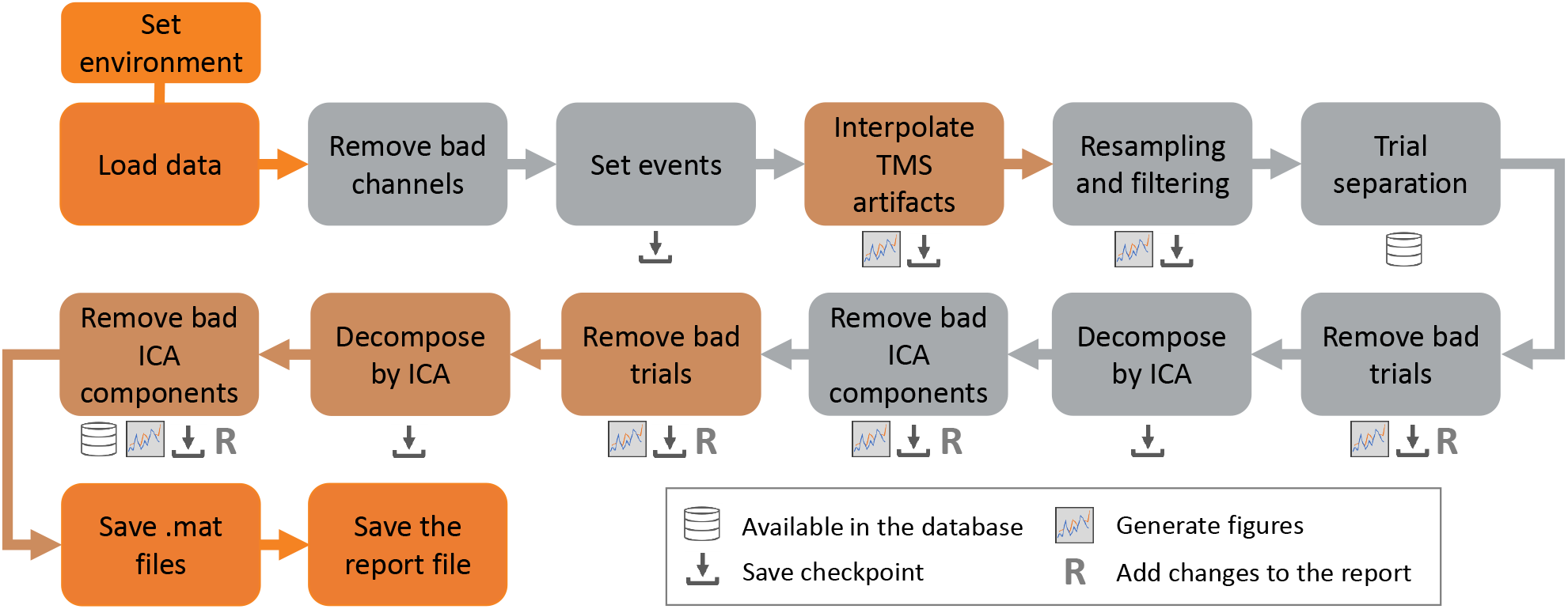
Code structure for processing the data. The seven fundamental sections are shown in gray, the optional sections in brown, and the extra functions for organizing the data in orange. The icons below each section indicate operations applied after the data processing step is complete.

After each data processing stage, the data were stored in specific folders that consist of `.set` files containing the pre-processing steps, a summary of the manual modifications made, and `.mat` files that document the event-related potential and spectral power density obtained for the analysis conducted. As a result, the outputs are organized and clearly annotated to ensure reproducibility and access to all the stages of the pipeline. For DDA, all data were utilized without filtering or downsampling.

#### Load data, remove bad channels, and set events

The pipeline was built to run one participant at a time. Initially, all the variables used to store the location of the necessary files were defined, then the folders for organizing the outputs were created, and the `.cnt` file was imported and saved as a `.set` file. Only the EEG recordings are of interest for this analysis, so the EOG and BIPs channels were removed. In addition, M1 and M2 were discarded. In total, 62 channels were processed, with CPz used as the reference and AFz as the ground electrode. Next, the database for each subject was updated with seven events for each phoneme pair task-related trial, namely: baseline, trial onset, first TMS pulse, second TMS pulse, first auditory stimulus, second auditory stimulus, and trial end.

#### Resampling and filtering

The data were resampled to 256Hz and two filters were applied: a notch filter with cutoff frequencies at 59Hz and 61Hz and a band pass filter with cutoff frequencies at 0.1Hz and 100Hz. The output generated by this last step is used for the subsequent analyses, differentiated only by the type of trial studied: (i) control, (ii) TMS applied to the lip target region, or (iii) with TMS applied to the tongue target region.

#### Trials separation

This section selects the analysis type and defines the first sound stimulus as the base event for ERP construction. This event sets an epoch separation that avoids undesired effects due to the high-amplitude TMS spikes.

#### Trials inspection and data cleaning

Two rounds of visual inspection were conducted on the data in search of trials with contaminated signals and channels with high-amplitude artifacts. To do so, the spectral power density (PSD) plots generated from the data were analyzed, in addition to the EEG recordings themselves. Using the EEGLAB interface, the unwanted segments were selected and removed from the analysis. After each round, ICA decomposition was applied using the library-adapted infomax ICA algorithm^65^. During ICA, 35 components were inspected and those that clearly exhibited an artifact signal were removed.

## Data Records

All of the data files may be accessed at the Open Science Framework repository ([OSF](https://osf.io/e82p9/)). Files are grouped by the year in which the dataset was collected: 2019 is labeled Study 1, and 2021 data is labeled Study 2. Each of these primary folders contains a second folder labeled “Data Descriptors”, and within each dataset folder are located dedicated subfolders for the raw data, processed data, and trial characteristics. The raw and processed data are grouped individually, with one subject per folder, and labeled according to Table 1.

### Raw and pre-processed EEG data

Raw EEG files were stored in the `.cnt` format. This format contains continuous EEG recordings saved over the EEG-TMS sessions. 66 channels were recorded, with electrode placement according to Figure 4. Pre-processed EEG data has also been made available in `.set` and `.mat` files, according to steps described in Figure 5.

### Event timestamps and behavioral data

For each trial, event timestamps are provided in `.csv` format, with one file for each recording session (Fig. 6). The events include (i) the second (final) TMS pulse of the pair, (ii) the sound stimulus onset, and (iii) the subsequent phoneme onsets. In addition to timestamps, the files provide labels for presented (true) and identified sound stimuli (phoneme or real/nonce word).

**Figure 6.**
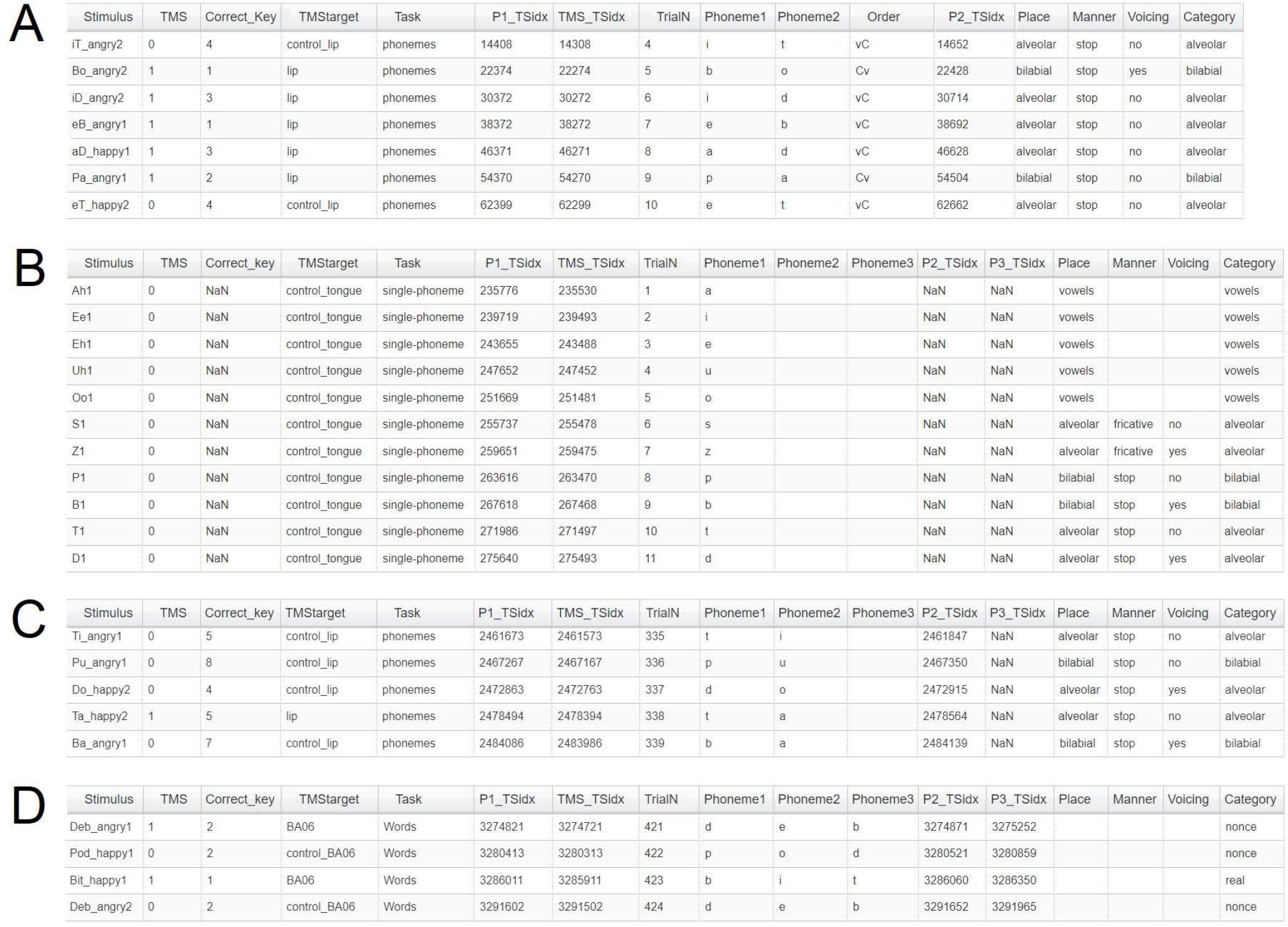
Files characterizing the EEG data. (A) The 2019 dataset provides labels for CV and VC phoneme pairs. The TMS condition in which control trials were collected is noted, as well as the articulatory features and category of each consonant phoneme per trial. A timestamp is provided for the onset of each phoneme and the final TMS pulse prior to stimulus presentation. Stimulation was conducted at only one cortical site per run. The control items are labeled according to the run in which they were collected. (B) The 2021 dataset provides the same information, when relevant, for single phonemes. (C) CV phoneme pairs are described in full, similar to the 2019 dataset. (D) Each component phoneme in the word trials is indicated and marked with a time stamp. Stimuli are categorized by word type.

### Technical Validation

Two sets of analyses were performed to support the technical quality of the datasets. Firstly, we extracted the grand mean event-related potentials (ERPs) by means of independent component analysis (ICA)^65^ to illustrate evidence of a stimulus-locked response across participants in each condition. We selected this method primarily due to its widespread use in the investigation of human cognitive information processing and therefore its familiarity among the electrophysiology research community. However, ICA can be subjective in its implementation by individual researchers, and the method may not be ideal for the analysis of specific types of data^66,67^. In particular, substantial attention has been paid to the need to remove the TMS artifact from TMS-EEG data^68–70^. Therefore, we performed a second analysis with delay differential analysis (DDA)^71–73^, a non-linear signal processing technique that requires minimal pre-processing and is noise insensitive^74,75^. The two analyses provide complementary evidence for the presence of a condition-dependent response in the EEG data. In particular, the DDA analysis illustrates excitatory activity during the time window of interest for our cognitive task, which differs by TMS condition.

#### Event-related potentials

The processing steps described in the Data Processing section were applied to the raw data to define the format of the auditory event-related potential (ERP) in the control condition and each TMS condition. The pipeline shown in Figure 5 was executed for all participants to ensure homogeneity in the analysis.

The mean and standard deviation of the potentials in the 1-second window after the first sound stimulus and the standard deviation for each of the 61 channels separated by the participant is represented in Figures 8 and 9 respectively. The reference pictures provided in Fig. 7 are meant to provide general guidance in interpreting the waveform; please refer to the cited papers for their original findings. We observe that the ERPs approximate the expected auditory-evoked potential (AEP) induced by phoneme pairs composed of stop consonants and vowels (see 7A, B). Deviations from the anticipated AEP may occur due to noise (our stimuli were immersed in white noise) and the exact combination of consonants and vowels in each stimulus item^76,77^. The shape of the TMS-evoked potential (TEP) will depend on the number of pulses delivered, the interpulse interval, and whether stimulation is subthreshold or suprathreshold. A wide variety of TMS paradigms have been tested with conflicting results, such that it may be better to observe the TEP in order to identify whether the paradigm was excitatory or inhibitory, or to consider the effect by means of an additional measure, such as a behavioral task (see 7C)^78^. The TMS paradigm used for collection of the dataset produced a facilitatory effect on performance in a phoneme perception task^51,54^.

**Figure 7.**
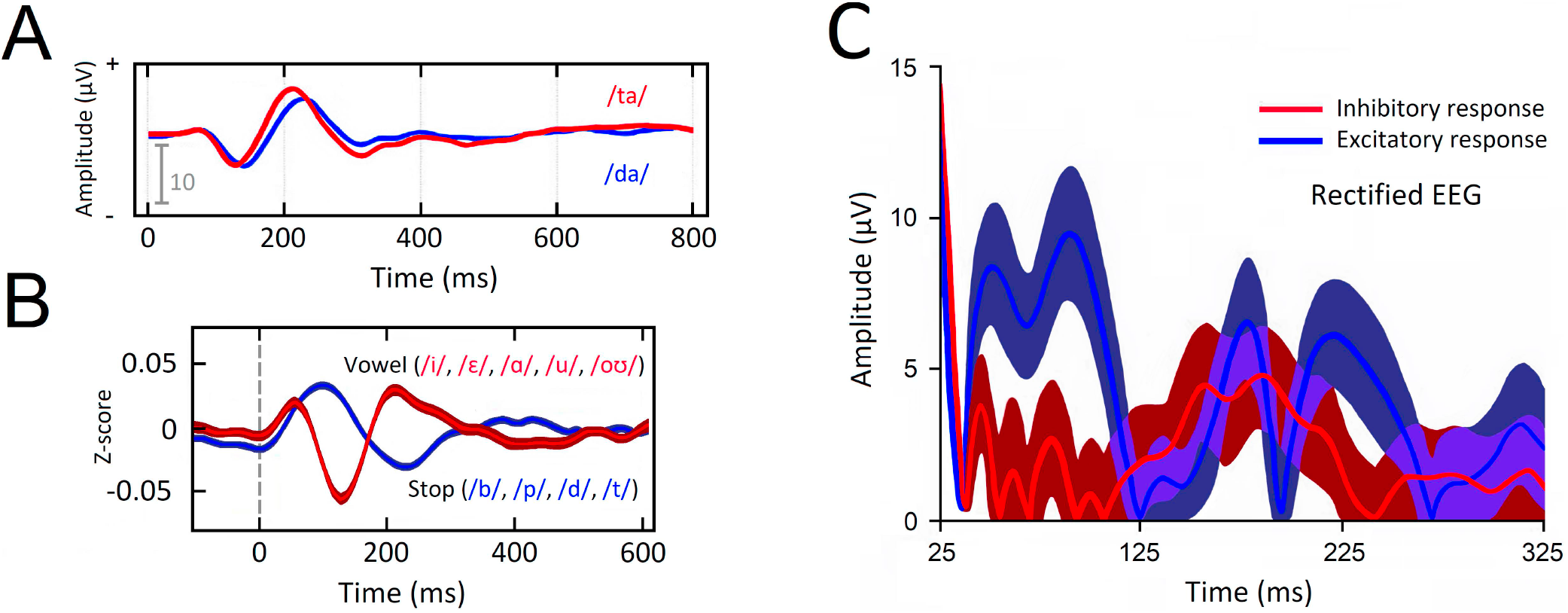
Expected waveforms. All plots were modified and reproduced with permission from the publishers. (A) The auditory-evoked potential (AEP) for stop (/d/, /t/) consonant-vowel pairs exhibits a small N100 potential followed by a larger P200 potential. Variation in timing will occur depending on the stimulus type and in the presence of noise^76^. (B) Vowels and consonants each produce a unique waveform^77^, such that the overall shape is dependent on the contribution of each to the waveform. (C) The shape of the TMS-evoked potential (TEP) will differ according to the cortical region targeted. Whether TMS creates an excitatory or inhibitory response can be observed in the shape of the resulting TEP. In the motor cortex, greater activity between 25-125 ms accompanies excitatory paradigms^78^. This figure illustrates the characteristic shape of the waveform for each type of neural response in the line and its standard deviation in the darker envelope.

**Figure 8.**
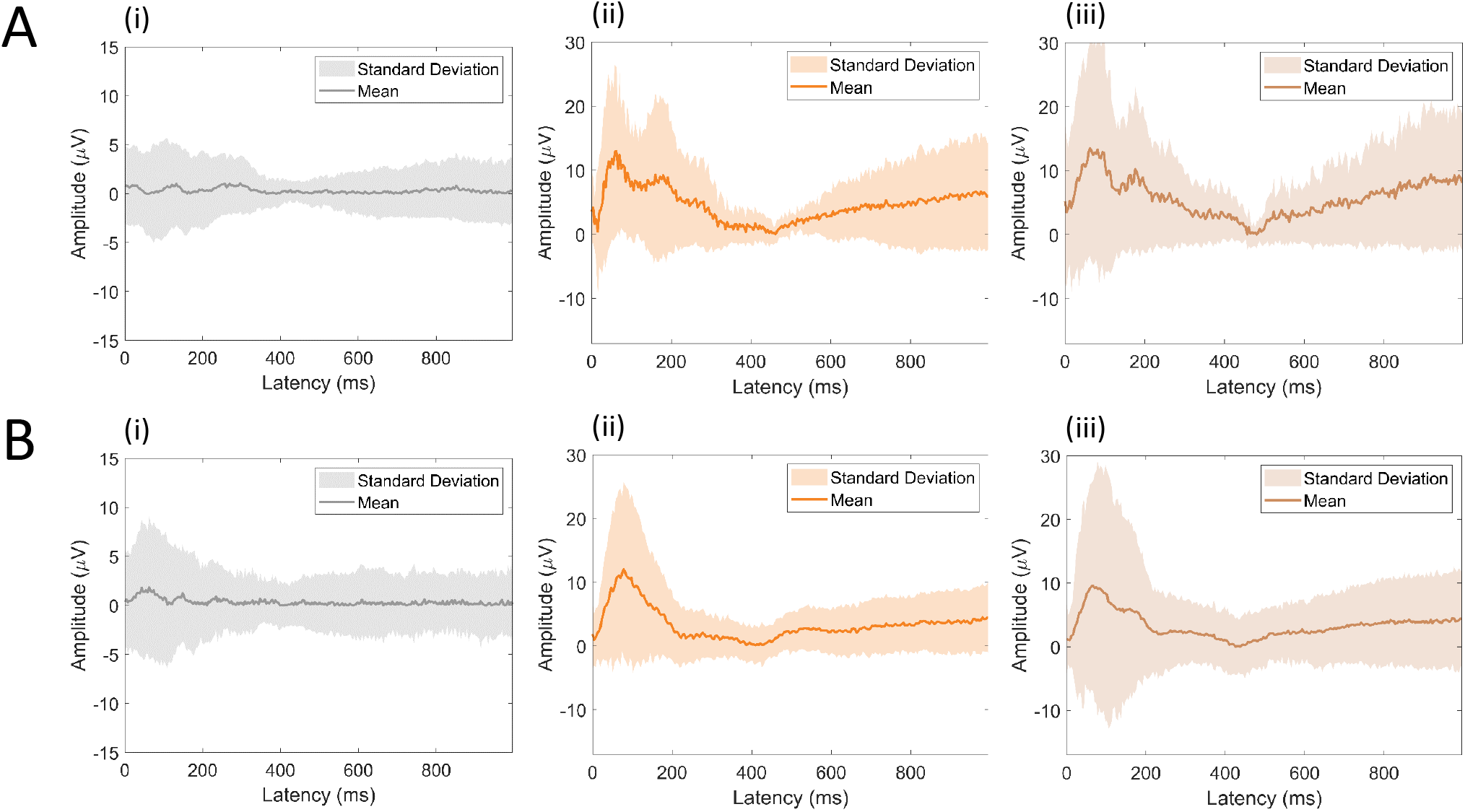
Event-related potentials (ERPs) from trial onset. The grand mean average ERPs to (i) control stimuli, (ii) stimuli with LipM1 stimulation, and (iii) stimuli with TongueM1 stimulation are displayed for (A) the 2019 and (B) 2021 datasets. The TMS conditions show rectified EEG activity to allow for comparison with the reference study.

**Figure 9.**
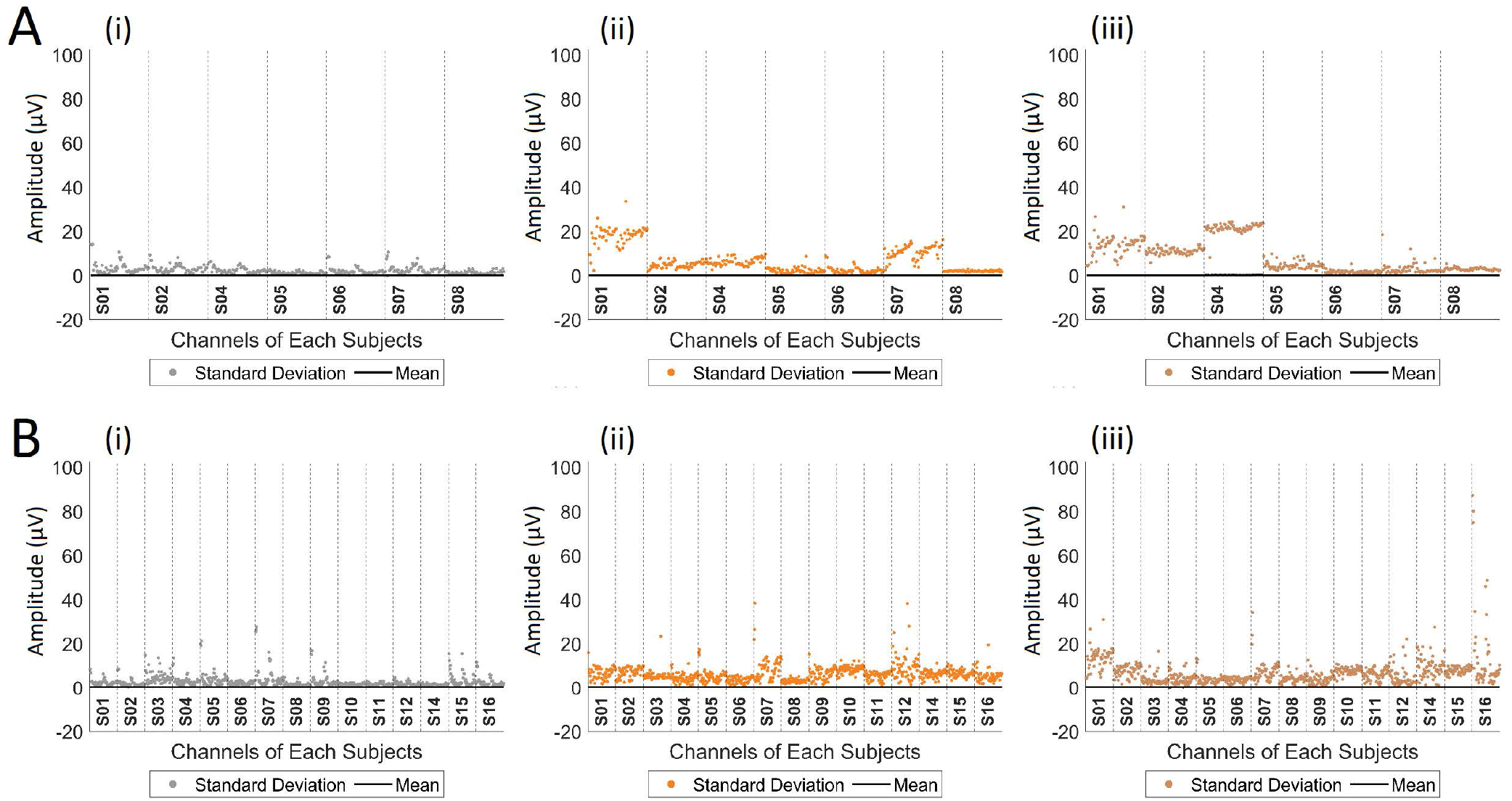
Characterization of the data by the mean and standard deviation of each channel per participant to (i) control stimuli, stimuli with LipM1 stimulation, and (iii) stimuli with TongueM1 stimulation are displayed for (A) the 2019 and (B) the 2021 datasets. The channels are arranged for each experimental subject from most anterior on the left to most posterior on the right.

Figure 9 provides an overview of the analyzed data in which each recorded channel is represented by its mean and standard deviation. Note that the value of the dispersion of the control trial group is smaller than the other values, as expected, and that some channels of certain subjects have high signal variation, especially those related to the frontal and medial portions of the brain, where stimulation occurred. This is unsurprising, given that TMS affects a unique subset of cortical neurons in each individual based on the position and orientation of neurons relative to the stimulation coil^79–84^.

#### Delay differential analysis

Delay differential analysis (DDA) is a signal processing technique that combines differential embeddings with linear and nonlinear nonuniform functional delay embeddings. The integration of nonlinear dynamics allows information from the data to be detected which may not be observable in traditional linear methods. DDA requires minimal pre-processing, which eliminates a highly subjective step in the data analysis. Sparse DDA models have several advantages over the high dimensional feature spaces of other signal processing techniques: (i) the risk of overfitting is greatly reduced; (ii) the sparse model concentrates on the overall dynamics of the system and cannot additionally model noise; (iii) DDA is computationally fast; (iv) there is no need of pre-processing except normalization to zero mean and unit variance for each data window in order to ignore amplitude information and concentrate on system dynamics. The DDA model consists of two sets of parameters: (i) the delays and model form are the fixed parameters that are kept constant throughout the analysis; (ii) the coefficients (*a*_1_, *a*_2_, *a*_3_) and the fitting error of the model are the free parameters. The coefficients are used as features to distinguish different dynamics in the data.

The DDA model used in this analysis is

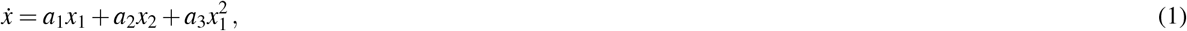

where *x_i_* = *x*(*t* − *τ_i_*). In this analysis, the fixed parameters are the same as in Ref^85^. We found that one of the free parameters, namely *a*_3_, can be used to describe neural activity in a manner similar to ERPs. However, an ERP and *a*_3_ are not strictly the same phenomenon. For details, see Ref^85^. Note that in most cases, there is no direct relation between frequencies and any of the model parameters, as explained in Ref^71^.

In the current analysis, the delays are *τ*_1_ = 6 *δt* and *τ*_2_ = 16 *δt*, with 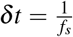, where the sampling rate is *f_s_* = 2000 Hz. These are double to the delays in Ref^85^ because the sampling rate is double. The window length is 30 ms and the window shift is 1 ms. In Fig. 10, we observe waveforms that display the same dynamics as the reference studies in Fig. 7. We observe neural activity 200 ms and 400 ms after stimulus-onset, the same time window where activity is observed in Fig. 7A. We also observe a sharp spike in activity 25-125 ms after the final TMS pulse, which corresponds to the results illustrated in Fig. 7B. This finding suggests excitatory activity.

**Figure 10.**
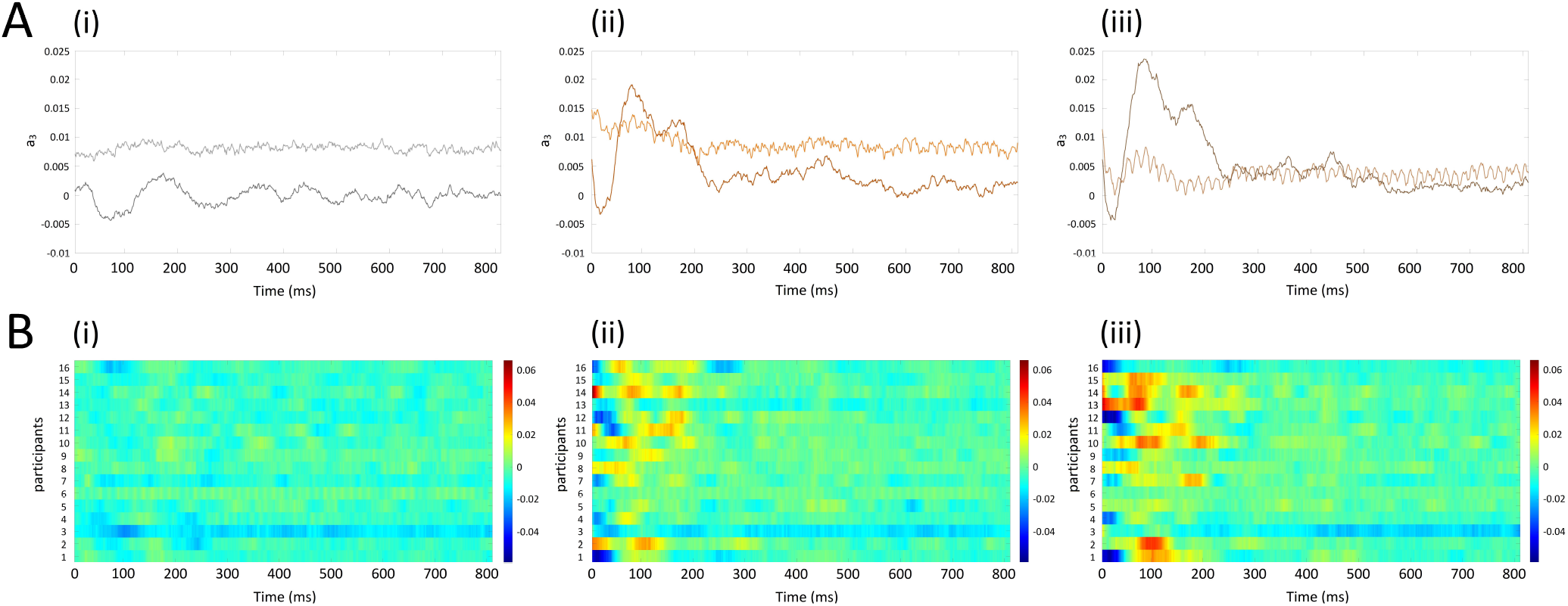
DDA coefficient *a*_3_ from Eq. (1) from trial onset. (A) The grand mean average DDA coefficient to (i) control stimuli, (ii) stimuli with LipM1 stimulation, and (iii) stimuli with TongueM1 stimulation are displayed. In each plot, the lighter line represents the 2019 data set, and the darker line represents the 2021 dataset. (B) The heatmaps for all individual participants are shown for the 2021 dataset.

#### Limitations and final remarks

This dataset is intended to provide researchers with a means to systematically test the classification accuracy of speech decoding models against naturalistic speech stimuli of increasing complexity, within and across datasets that manipulate the cortical state of participants. To our knowledge, this is the first EEG dataset for neural speech decoding that (i) augments neural activity by means of neuromodulation and (ii) provides stimulus categories constructed in accordance with principles of phoneme articulation and coarticulation. Nonetheless, several limitations of the dataset can be noted.

First of all, the experimental task involves aspects of comprehension, production, and motor activity (in the form of a button-press response), which may be subject to some overlap in the neural signal. In particular, in single phoneme trials, speech may have been produced while potentials relevant to comprehension of the speech sound were still ongoing. However, it is well known that the neural networks underlying motor and language functions are not strictly dissociable. They are frequently coactivated, even in covert speech or comprehension paradigms^45,46^. Therefore, we believe this phenomenon underlies most if not all speech decoding paradigms, to a greater or lesser degree, and likely represents neural processing in a naturalistic context.

Secondly, inner speech is widely adopted in the speech decoding literature, where it is often considered to be the most intuitive way of controlling a BCI device. However, inner speech decoding paradigms may not accurately mark the onset of individual stimuli or component phonemes in the recorded data. We believe that prior to transitioning to an inner speech paradigm, researchers would benefit from developing models that can target specific features of the speech stream, such as articulatory features and coarticulation. This kind of systematic study of the speech input may lead to more robust models overall and a better understanding of how this process occurs, rather “black box” models that must be trained on huge amounts of data or rely more heavily on predictive language models than on actual decoding accuracy. Both of these trends in the literature address issues that are orthogonal to the improvement of the actual decoding model.

Questions may also arise as to how neuromodulation may be integrated into a BCI device. We believe that greater attention is needed to study the possibilities for applying neuromodulation during speech decoding. At this time, there is no viable BCI device for evoked paradigms that can be used by the target user population, even with an inner speech paradigm, and therefore any talk of a full-functional device that is ready for end users would be premature. As the study of this procedure continues, we may find a means to utilize neuromodulation for model or participant training, and new possibilities for fast and accurate neuromodulation techniques that could be integrated into a headset continue to be developed.

Finally, we have noted that the two datasets differ in size and quality. We recommend that the larger dataset be used for model development and that the smaller dataset be used to validate the model in more stringent conditions. Our own tests have shown that, by means of DDA, we can successfully perform a classification analysis with both data sets^51^.

## Usage Notes

The raw .cnt EEG files can be read in MATLAB with the FieldTrip Toolbox^86^ and in the Brainstorm^87^ eepv4_read.m function, or in Python with the libeep library. The pre-processed files can be read in MATLAB with EEGLab^64^, FieldTrip^86^, or in Python with MNE^88^.

## Code availability

The data and codes used in this work are available at [OSF](https://osf.io/e82p9/) to allow reproducibility and sharing of information under the CC-BY-NC-ND 4.0 license (http://creativecommons.org/licenses/by-nc-nd/4.0/). The routines can be found in the Study/EEG_Data_Processing/Code folder. These routines are responsible for the analyses presented in the technical validation section. The results obtained for both signal processing techniques, as discussed in Data Processing section, are placed in the same folder. The same code is available on [GitHub](https://github.com/mcjpedro/speech_decoding) so as to allow for version control and discussion of the implementation and analysis carried out in this work. The routines were built to obtain the ERP using only ICA and signal cleaning was performed using the pipeline described in Figure 1, based on the EEGLab library versions 2022.0 and 2022.1 native to MATLAB.

## Author contributions statement

L.C. conceived the experiments, ran the experiments, created figures, and wrote the manuscript. J.P.C.M. developed codes, analyzed the results (ICA), created figures, and wrote the manuscript. V.R.C developed codes, analyzed the results, and wrote the manuscript. C.L. analyzed the results (DDA), created figures, provided technical feedback, and reviewed the manuscript. E.M.A.M.M., A.F., and T.J.S provided technical feedback and reviewed the manuscript.

